# Peas on Mars: A study of garden pea, *Lathyrus oleraceus* (=*Pisum sativum*), growth in Martian regolith simulant treated with black soldier fly frass

**DOI:** 10.1101/2025.11.14.688485

**Authors:** J. Emmanuel Mendoza, N. B. Lemke, J. K. Tomberlin

**Author notes:** Credit AUTHOR ROLES JEM - Conceptualization, Methodology, Software, Validation, Formal Analysis, Investigation, Resources, Writing-Original Draft, Writing-Review & Editing, Visualization; NBL - Conceptualization, Methodology, Software, Validation, Formal Analysis, Data Curation, Writing-Original Draft, Writing-Review & Editing, Visualization, Supervision, Project Administration; JKT - Methodology, Resources, Writing-Review & Editing, Supervision, Project Administration, Funding Acquisition.

## Abstract

Future long-duration missions to Mars will require *in situ* food production systems to reduce dependence on resupply missions and enhance crew autonomy. This study evaluated the growth performance of peas, *Pisum sativum,* cultivated in Martian regolith simulant (MMS-1) ameliorated with black soldier fly, *Hermetia illucens*, (BSF) frass at inclusion levels of 0, 10, 25, and 50% by volume. Pea plant fitness was then assessed over the course of two, 8-week greenhouse trials by measuring germination rate, plant height, leaf chlorophyll content, and final dry biomass. These were compared against peas grown in a control substrate consisting of commercially available gardening soil treated with the same frass-inclusion percentages (0, 10, 25, or 50% by volume). Germination and height responses were statistically similar between regolith and soil, suggesting that properly amended regolith may be able to sustain *in situ* crop growth on Mars. Notably, peas grown in regolith simulant + frass exhibited slightly higher chlorophyll content than the controls, while biomass productivity was lower than commercial potting mix + frass when comparing across the same frass-inclusion percentage. These findings represent the first experimental evidence that BSF frass can enhance crop cultivation in a Martian soil analog, highlighting its potential role in bioregenerative life support systems and in-situ resource utilization (ISRU) strategies. In addition, fitting quadratic regression models to the data indicated an optimal frass inclusion between approximately 5–32% (depending on which metric was to be maximized), beyond which fitness declined. Beyond space applications, these findings build on prior studies by further supporting the value of insect-derived amendments for improving nutrient-poor or degraded soils.

Table 1:
Glossary of Acronyms and Initializations Used

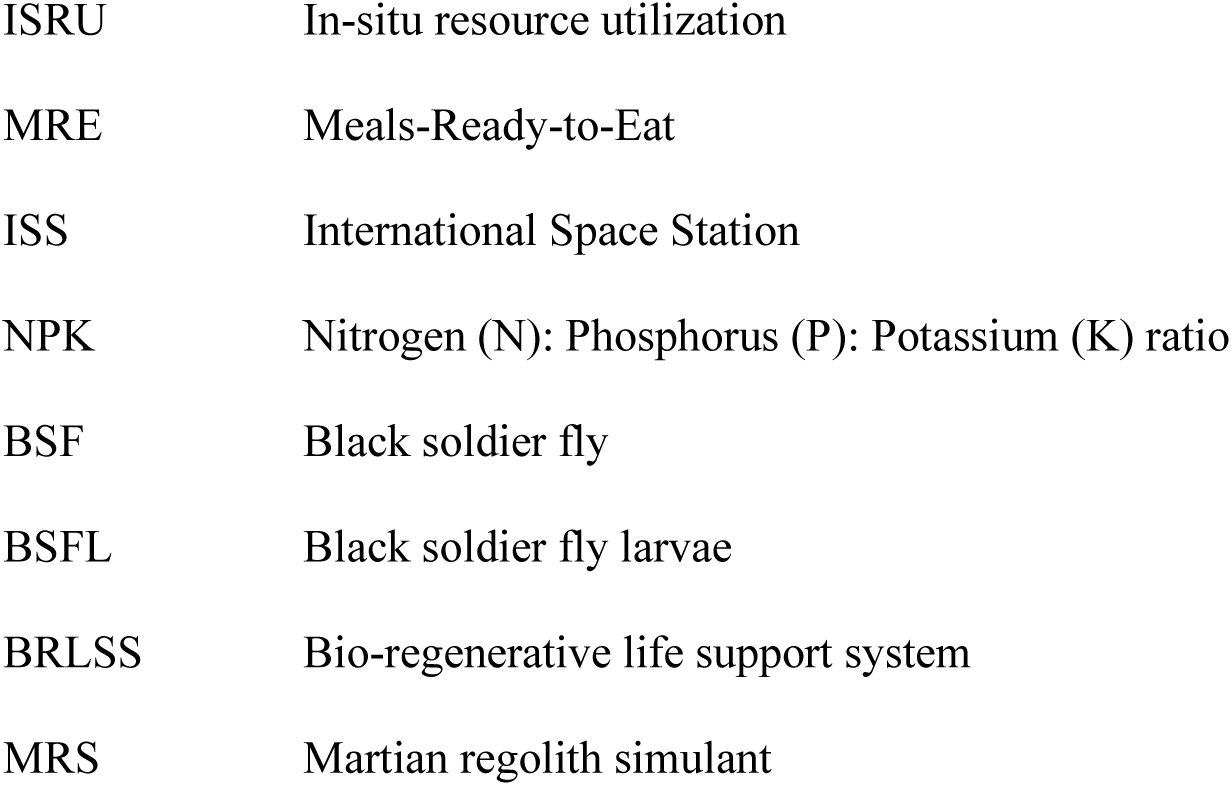

## 1. INTRODUCTION

### 1.1 Astrobiology

Space technology development has historically focused on rockets and materials science. However, now that many space programs are actively planning manned missions to the lunar surface and Mars, advances in astrobiology and space agriculture will be necessary for astronauts to survive on long-term missions [1], especially considering that the shortest feasible human-crewed Mars mission would likely last no less than 15 months [2]. More likely crewed missions to Mars will approach 34 months to allow for extended surface operations, habitat setup, and in-situ resource utilization (ISRU) development [3], which begs what will astronauts eat during those 15-35 months.

Currently, the food provided to astronauts on manned missions consists of heat-stabilized pre-packaged meals, e.g., Meals-Ready-to-Eat (MREs); however, because there is a hard threshold of cargo that can be brought aboard manned space missions (both by weight and volume), an important trade-off exists between the weight of bringing aboard additional food with the weight of water and oxygen needed for life support systems [4]. Of course, re-supply missions might be uncrewed, but delays or failures might jeopardize the survival of crews in space [5]. As such, finding methods of growing food-plants on space stations and on extraterrestrial surfaces (Moon, Mars) will be a crucial breakthrough for ensuring astronauts have access to adequate calories and nutrition throughout multi-year missions [6].

Studies conducted on the International Space Station (ISS) have demonstrated that leafy vegetables like red romaine lettuce, *Lactuca sativa* cv. ‘Outredgeous’, can be safely grown and consumed by astronauts, offering a fresh food source to supplement conventional MREs [7]. Additionally, microgreens such as green daikon radish, *Raphanus sativus var. longipinnatus*, and rioja radish, *Raphanus sativus ‘Rioja’,* have been identified as promising crops for space missions due to their fast growth and high levels of bioactive compounds (e.g., vitamins, antioxidants, minerals, and phytonutrients) [8]. The literature contains several studies focused on growing crops in Lunar Regolith [9,10]. However, to date, relatively little *ex situ* research has been conducted on growing crops on Mars, and so this study addresses this knowledge gap.

### 1.2 Martian Regolith

Unlike Earth, the surface of Mars is covered with regolith rather than soil [11]. Martian regolith contains some of the essential nutrients for life, such as potassium (K), calcium (Ca), magnesium (Mg), and iron (Fe). These elements primarily occur in the forms of potassium oxide (K2O, calcium oxide (CaO), magnesium oxide (MgO), and both ferrous iron (II) oxide (FeO) and ferric iron (III) oxides (Fe2O3), along with minerals like plagioclase (Na₁₋ₓCaₓAl₁₊ₓSi₃₋ₓO₈), zeolite (Mx[(AlO2)x(SiO2)y]·nH2O), and hematite (Fe2O3) [12]. But by contrast to soil, regolith chiefly lacks organic matter, microscopic/minute forms of life (e.g., bacteria, archaea, algae, fungi, protozoa, gregarines, nematodes, springtails, etc.) [13], and some macronutrients like Nitrogen (N) and phosphorus (P) [14]. Additionally, the presence of toxic perchlorates (e.g., calcium perchlorate (Ca(ClO4)2), magnesium perchlorate (Mg(ClO4)2)) in Martian regolith poses a major obstacle to life, as they inhibit plant growth, even after partial removal [15]. Furthermore, water on Mars is present in the form of highly-saline brine and ice, and will likewise require specialized detoxification techniques to make water suitable for irrigation [16]. As such, it is theorized that Martian regolith can potentially be used as a growing medium for crops, but its composition presents significant challenges that must first be ameliorated [17]. This experiment focuses on addressing one component of this complex task.

### 1.3 Insect Frass

Utilizing insect frass derived from mass-reared insect colonies is one potential way to enhance Martian regolith’s deficiencies in nitrogen, phosphorus, and organic matter. For instance, larvae of the black soldier fly (BSFL), *Hermetia illucens*, (L.) (Diptera: Stratiomyidae), are mass-reared throughout the globe for waste remediation [18,19]. As such, an industrial BSF-rearing system is designed to convert sometimes up to 100 tons of agricultural waste per day into insect biomass (i.e., as whole, pureed, or powdered larvae) to then be used as animal feed. However, roughly ∼60% of the organic waste is converted into frass as a byproduct. Technically ‘frass’ refers only to the powdery waste excreted by the larvae [20], but more commonly the word is used to describe the residual material remaining after a feedstock is digested by BSF larvae (and so is actually the mixture of both frass and undigested feedstock). Since frass (as defined this way) represents the largest output of a mass-rearing operation, a great deal of research has focused on applications for BSFL frass in terrestrial agriculture and found many positive benefits, including high levels of N, K, and microbes (see section 1.4 for details). For example, a one-month composting trial showed that Martian regolith simulant amended with BSFL had far higher plant-available nutrients than untreated regolith simulant, with increases of +1179% in potassium, +803% in phosphorus, +67% in magnesium, +7847% in manganese, and +188% in nitrogen, from 19.8 ppm to 57.1 ppm [21].

Of course, one might question whether insects can survive and perform well aboard space missions; the common fruit fly, *Drosophila melanogaster* Meigen (Diptera: Drosophilidae), has a long history of being studied aboard the ISS, with investigations focusing on genetic, developmental, and immune responses to microgravity and radiation [22–24]. As such, *D. melanogaster* serves as a comparative model for evaluating the potential of other insects in space, such as BSFL. Indeed, research has now shown that BSFL can survive high-G forces, akin to those seen during launch [21] and BSFL can also successfully pupate and develop on Martian regolith simulants [21]. Together, these results suggest the possibility that BSFL can be cultured in space, and that their frass can be used as an *in situ* soil enhancer in terraforming projects on the Moon and Mars [21,25–27]. Though still speculatory at this point, an idealized bioregenerative life support system (BRLSS) could thus utilize BSFL to upcycle organic wastes (e.g., astronaut urine and feces, inedible parts of plants, food discard) produced during deep space missions and off-world settlements into BSFL biomass (for direct consumption or as animal feed) and frass. Such a concept is not so far-fetched, since (a) BSFL can digest and develop on a wide range of organic wastes, (b) insect protein is eaten by many cultures throughout the world [cite], (c) as astronauts aboard the ISS already drink water that is filtered from their own urine [28]. Indeed, futurists have also increasingly realized that insects will play a role in food for humans and animals on Earth and in space [29], and other candidates besides BSF include the yellow Mealworm, *Tenebrio molitor* L. (Coleoptera: Tenebrionidae), the domestic cricket, *Acheta domestica* (L.) (Orthoptera: Gryllidae), among others.

### 1.4 Frass as a soil amendment

Insect frass is nutrient-dense and also has a relatively-even ratio of phosphorus pentoxide (P₂O₅) to potassium monoxide (K₂O) (i.e., NPK ratio), making it similar in composition to commercial organic fertilizer [30,31]. Moreover, it is highly microbially active, and because of these qualities, the amount of research articles studying applications of frass (e.g., soil amendments, pest control, animal feed) has recently exploded [32,33]. One such study found that BSFL frass enhanced the growth of maize, *Zea mays,* and the highest application rate of frass (of 7.5 metric tons ha^-1^) increasing yields by 14% above commercial organic fertilizer [34]. Similarly, another study found that when frass was given to pak choi, *Brassica rapa subsp. Chinensis*, significantly higher plant biomass was observed compared to the chemical fertilizer control group [20]. While there exists plenty of research to indicate the beneficial qualities of using BSFL frass to improve terrestrial crop production, none yet exists that evaluates the potential for growing plants in Martian regolith. While a prior experiment found that using commercial horse/swine manure supplementation markedly improved viability and productivity of plants grown in both Lunar and Martian regolith simulants [35], rearing horses and pigs in space is obviously less feasible than insects, and so within the context of *in situ* resource utilization, insect frass warrants exploration.

### 1.5 Research Objectives

To address this gap, the objective of this study was to compare the effect on the plant growth of common garden peas, *Lathyrus oleraceus = Pisum sativum,* of (a) Martian regolith simulant supplemented with BSFL frass to (b) commercial gardening soil also supplemented with BSFL frass. The level of frass included in both regolith and commercial gardening soil was varied along a gradient of 0-50% frass, measured by volume. Peas were selected because as legumes they form symbioses with nitrogen-fixing bacteria, and so the soil did not need to be inoculated. Moreover, research has shown that frass can be slightly alkaline, which garden peas perform well in (USDA.gov). Pea plant fitness was assessed by measuring germination rates, plant height, leaf chlorophyll content, and final dry biomass. We assumed a null hypothesis that Pea fitness would be equivalent regardless of whether peas were grown in Martian regolith simulant or commercial gardening soil, when comparing metrics across identical frass supplementation levels.

## 2. METHODS

Experiments were conducted at the F.L.I.E.S. Facility (Texas A&M University, College Station, Texas, USA). Attached to the facility is a greenhouse measuring 23.4 m by 9.3 m, and its interior is divided into three sections. The study was conducted in the southwestern-most of these sections. Each greenhouse section was equipped with standard rolling tables, a wet wall for evaporative cooling, as well as industrial fans to remove hot air. Despite these features, extreme temperatures occasionally occurred due to the recently-replaced glass windows which were not yet treated with shade paint. The southwestern greenhouse was also home to an active BSF colony, and although some adult escapees were occasionally found perching on the plants, their presence did not otherwise interfere with the outcomes of the study.

Two trials were conducted: Trial A (occurring from September 8, 2023 - November 10, 2023) and Trial B (occurring from September 15, 2023 - November 17, 2023), each with a duration of approximately ∼60 days that corresponded to the average planting-to-harvest time for garden peas. Since the date ranges of Trial A and Trial B overlapped, the two were set-up on opposite sides of the greenhouse (with Trial A set up on the right-middle table of the greenhouse, and Trial B set up on the left-middle table). Each trial was terminated when plants had fruited and began to wither.

### 2.1 Frass Acquisition

BSFL frass was sourced from a local producer (EVO Conversion Systems LLC, Bryan, TX, USA) by mechanically sifting the larvae from the remaining material at the end of their feed cycle. This producer fed their larvae on a combination of spent brewer’s mash and leftover bread sourced from local restaurants. Prior to each trial, frass sat unmodified for 7 days within an opaque plastic garbage bag stored in walk-in incubator “Chamber 2” at the FLIES facility, which was maintained at 26°C and 60% RH, with 14:10 L:D cycle.

### 2.2 Experimental Design

Each experimental unit consisted of a 1.89-L black plant nursery pot (Oubest, Fuzhou, China) (Height = 0.13-m, upper D = 0.15-m, lower D = 0.105-m), filled with substrate (see section 2.3). Each round pot had drainage holes to prevent waterlogging. Into each of these, n = 2 non-GMO Lincoln variety garden pea seeds (Harris Seeds / Garden Trends, Rochester, NY, USA) were planted at a depth of 1-2 cm beneath the substrate surface, following conventional methods (USDA.gov). We decided to use two pea seeds / unit to double the chance of at least one viable plant growing per pot, but also give each plant sufficient space if all (i.e., both) seeds germinated.

### 2.3 Substrate

To test the effect of frass inclusion on pea growth, treatments varied the amount of frass along a continuum with four levels: 0%, 10%, 25%, and 50%, which were measured by-volume (as calculated based on volumetric displacement). These amounts were added either to Mohave Mars Regolith Simulant MMS-1 (The Martian Garden, Austin, TX) or “Kellogg’s All Natural Garden Soil” (Kellogg Garden Products, Lockeford, CA) that acted as a control. Preliminary experimentation showed that pea plants did not grow when frass constituted more than 50% of the growth media, which was consistent with other studies [36]. Because of this, only proportions of frass below 50% were tested. Each trial had a total of 48 experimental units, given n = 4 frass-inclusion percentages, n = 2 types of substrates, replicated n = 6 times. The growth media for each treatment was prepared by mixing frass with either commercial gardening soil or regolith simulant in a single Sterilite container (33.0 cm L ⨉ 21.6 cm W ⨉ 30.5 cm H), and is a standard method to ensure that the substrate was relatively homogeneous, rather than forming distinct layers of frass on top of regolith/soil [37]. The total amount of substrate and frass needed for each respective treatment level was weighed using a digital kitchen scale (BAGAIL BASICS Digital Kitchen Scale, BAGAIL, Suzhou City, China) and then divided evenly among the six replicate nursery pots.

### 2.4 Experiment Set-Up

Experimental units (i.e., nursery pots containing n = 2 planted Peas) were placed atop a rolling table in one of eight, 2 ⨉ 3 arrays, then randomized. To prevent water runoff, paper plates (21.6 cm D, Dixie brand, Georgia-Pacific Atlanta, GA) were placed underneath each pot as improvised plant saucers. To protect the peas from too much sun exposure, 40% shade cloth (Kesfitt Patio Direct, Guangzhou, China) was suspended above the table at a height of 0.76 m. To complete the set-up, all experimental units were watered with 50 mL or reverse osmosis (RO) water. Photos of the set-up can be found in Figure S1. Regular watering continued on day 2 for Trial A and day 3 for Trial B, (which for logistical reasons ensured both trials were watered on the same weekday, reducing the burden of labor). From this point forward, each pot was watered every two days, by providing RO water *ad libitum* from a graduated beaker between 1200-1300 h to fully soak the substrate (approximately 50 mL). After four weeks, a single poplar dowel (0.635 cm diameter, 1.22 m height; Madison Mill, Ashland City, TN) was inserted vertically into the soil of each pot to promote vertical growth, as peas typically require a trellis for support (TAMU Extension).

As a precaution (i.e., in case of abnormal weather conditions), ambient environmental conditions – viz., dry-bulb temperature (℃), relative humidity (%RH), and light intensity (lux) – were automatically recorded every 60 minutes using a HOBO Temp/RH/Light/Ext-Analog Data Logger MX1104 (Onset Brands, Bourne, MA); though this data was not considered as part of our statistical analysis.

### 2.5 Data Collection Schedule

Data were collected weekly between 13:00 and 16:00 h (prior to watering) for 8 weeks. As proxies for fitness, the germination rate per pot (completed during the first week), plant height, and leaf chlorophyll content were all measured each week. At the end of the experiment, the dry biomass of each experimental unit was calculated, and any harvested fruit was also weighed.

Germination rate (quantified as: no peas germinated = 0%, a single pea germinated = 50%, both peas germinated = 100%) was based on a visual inspection of whether the first clamshell leaf had appeared from the soil (i.e., the thick, rounded cotyledons that emerge from the seed during early growth). Plant height was considered as the distance between the soil surface and the farthest extent of the plant’s above-ground structure (whether it was a leaf or tendril). This distance was measured for both plants within each experimental unit using a metal ruler. During this process, pea plants were manually manipulated to ensure the stem was extended to its maximum length. Height represents the maximum value recorded per experimental unit rather than the mean of both plants. Height data were collected throughout the experiment, but only end-of-study measurements were analyzed. Chlorophyll content was measured by haphazardly choosing a matured leaf and clamping down on the largest cross-sectional area with a SPAD 502 Plus Chlorophyll Meter (Spectrum Technologies, Inc., Plainfield, IL, USA). Chlorophyll measurements reflect the overall leaf greenness and plant health, with higher SPAD values indicating greater photosynthetic capacity. Dry biomass was determined as the total oven-dried weight of the entire plant, not the peas themselves, and values were normalized by the number of plants present within each experimental unit to account for variation in survival and germination. Generational effects were not established in this study, so all results reflect single-generation responses to frass inclusion and substrate type.

To assess dry biomass at the end of each trial, plants were first gently uprooted from their pots, and stuck soil was removed by washing the plant roots in RO water, taking care to preserve the root ball. Approximately 50-100 g of the substrate still remaining in the pots was collected from each replicate and preserved in a Kenmore chest freezer (Kenmore, Chicago, IL, USA) in the back building of the FLIES Facility for future analyses. The washed plants were then individually placed into paper bags and also stored in the same freezer as the substrate samples. The plants were dried in a Despatch Countertop Medical Oven (Despatch, Lakeville, MN, USA) at 55 ℃, until the weight loss from evaporation plateaued after 24 hours (< 1% mass change). Plants were then weighed using the A&D ER-182A Electronic Balancing Scale (A&D Weighing, Ann Arbor, MI, USA) to yield dry biomass.

### 2.6 Statistical Analysis

Data were cleaned in MS Excel (version 2410), and exported to Python (version 3.8.8) for statistical analyses. Data was plotted utilizing standard libraries (e.g., pandas, matplotlib, scipy.stats, scikit_posthocs, and statsmodels). Preliminary analyses revealed that the data (plant height, dry weight of peas, chlorophyll content) were non-parametric, as indicated by a visual inspection as well as a significant Shapiro-Wilk test (p<0.05). Therefore, non-parametric statistical analyses were performed. The Mann-Whitney *U* (a.k.a., Wilcoxon Rank-Sum) was used when comparing only 2 groups (e.g., when comparing grouped treatments between regolith and the soil control). These tests used an alpha-level of 0.05 (prior to any p-adjustment), and (except for germination rate) all zeros instances were ignored for statistical tests since it did not make sense to consider the height/chlorophyll/weight of plants that did not grow. To supplement these analyses, regression models were fitted to the data to examine the effect of increasing frass supplementation on each of our metrics. Ultimately, quadratic regression models were selected for all response variables, as they consistently provided a better fit to the data based on higher R² values (Table 2), and/or because these functions had a single critical point that predicted where the optimum frass inclusion level was.

**Table 2.**
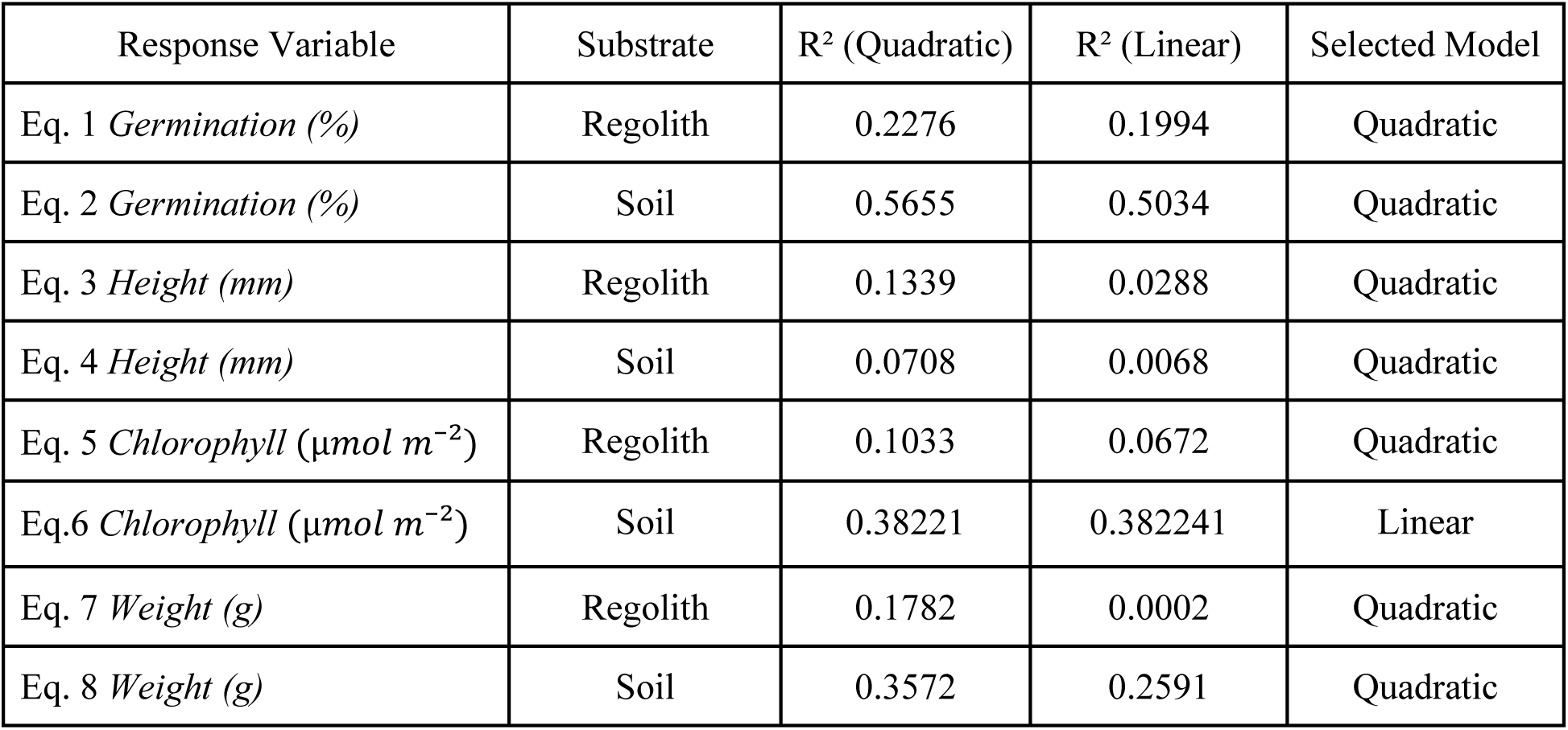
Comparison of R2 coefficients for linear and quadratic regressions used in model selection. For each substrate response variable, a linear and quadratic regression model was developed, and the model with the highest R2 was selected.

## 3. RESULTS

The objective of this study was to compare the effect of simulated Martian regolith supplemented with BSFL frass to a control of commercial gardening soil also supplemented with BSFL frass on the growth and fitness of common garden peas. Mean values for fitness are expressed in Table 3.

**Table 3.** Mean values of fitness metrics for each treatment. Means are given followed by ± standard deviation. Zeros are included for Germination (%), but have been removed for Height, Chlorophyll, and Biomass. * Indicates statistically different groups when comparing Regolith vs. Soil of the same Frass Inclusion (%) using a Mann Whitney *U* test.

| Frass | Medium | Height (mm) $\mu \pm$ | Biomass (g) $\mu$ | Germination (%) | Chlorophyll |
| --- | --- | --- | --- | --- | --- |

| Inclusion (%) | | $\sigma$ | $\pm \sigma$ | $\mu \pm \sigma$ | $(\mu\text{mol}/\text{m}^2) \mu \pm \sigma$ |
| --- | --- | --- | --- | --- | --- |
| 0% | Regolith | $180.23 \pm 57.77^*$ | $0.199 \pm 0.15^*$ | $70.83 \pm 39.65$ | $27.76 \pm 7.09^*$ |
| 0% | Soil | $146.6 \pm 20.75^*$ | $0.341 \pm 0.18^*$ | $91.67 \pm 19.46$ | $17.49 \pm 7.81^*$ |
| 10% | Regolith | $193.16 \pm 80.16$ | $0.386 \pm 0.24^*$ | $79.17 \pm 33.43$ | $29.30 \pm 10.28^*$ |
| 10% | Soil | $167.47 \pm 41.46$ | $0.722 \pm 0.41^*$ | $87.50 \pm 22.61$ | $23.73 \pm 7.15^*$ |
| 25% | Regolith | $225.00 \pm 49.95$ | $0.384 \pm 0.27$ | $62.50 \pm 43.30$ | $36.53 \pm 2.24$ |
| 25% | Soil | $156.44 \pm 26.47$ | $0.669 \pm 0.42$ | $79.17 \pm 33.43$ | $32.82 \pm 8.29$ |
| 50% | Regolith | $123.00 \pm 40.83^*$ | $0.197 \pm 0.12^*$ | $25.00 \pm 39.89$ | $32.84 \pm 9.03^*$ |
| 50% | Soil | --- | --- | $16.67 \pm 32.57$ | --- |

### 3.1 Germination Rates

Germination rates were first compared between peas planted in regolith + frass and the soil + frass control. Data were treated as ordinal percentages (0%, 50%, and 100%) based on whether neither, one, or both, seeds within each replicate germinated. Most interestingly, a majority of peas (70.83%) germinated in Regolith without any added frass (Table 3). In most cases the mean germination rate was higher for peas planted in Soil + Frass, except at inclusion levels of 50%, since no peas germinated in soil with 50% frass. When pooling germination rates across each substrate type, a Wilcoxon rank-sum test indicated no significant overall difference between germination rates of peas planted in regolith + frass compared to soil + frass (p = 0.313) (Figure 1), and indeed pairwise comparisons were all statistically insignificant (Table 3). To determine optimal frass inclusion levels for each substrate type, quadratic regressions were fit to the data. For peas planted in regolith + frass, only the intercept (*p = 9.47E-09*) was found to be significant, suggesting co-linearity between the terms or some other form of non-linearity (Eq. 1). For peas planted in soil frass, both the intercept (*p = 6.1E-6*) and quadratic (*p = 0.0147*) terms were found to be significant, with the linear term (*p = 0.597*) being highly insignificant, suggesting a curvilinear response (Eq. 2). The local maxima for each were found to be 6.9% and 5.6% for regolith and soil, respectively.

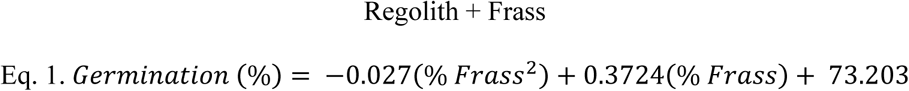

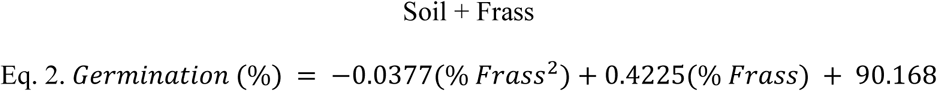

**Figure 1.**
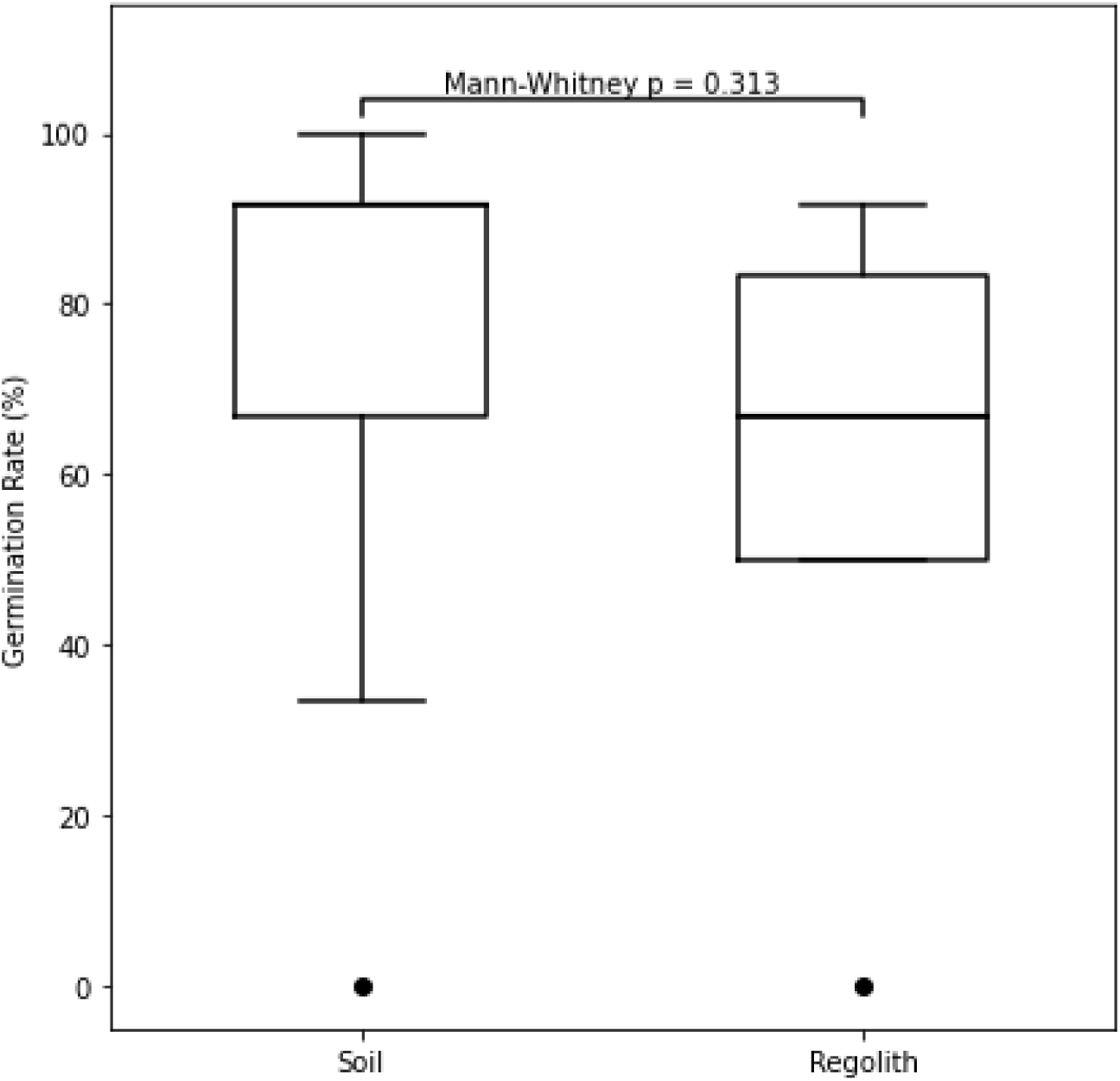
Boxplot comparing germination rate (%) between *Pisum sativum* grown in soil and regolith simulant. Each experimental unit consisted of two pea (*Pisum sativum*) seeds planted in 0.13-m tall, 0.15-m upper length & depth, and 0.105-m lower length & depth square pots filled with substrate, of which 0%, 10%, 25%, or 50% was BSF frass by volume. Mann-Whitney U test (ɑ = 0.05) was used to compare differences across treatments pooled by frass-inclusion percentage.

### 3.2 Plant Height

Although mean plant height was in all cases taller for peas planted in regolith + frass versus those planted in soil + frass (Table 3), a Wilcoxon Rank-Sum test detected no significant difference (*p = 0.68*) between peas planted in regolith or soil when all frass-inclusion levels were pooled for each substrate type. When comparing plant height in response to substrate type for the same levels of frass, Regolith and Frass were statistically different for 0% frass inclusion, as well as 50% frass inclusion (Table 3).

To determine optimal frass inclusion levels for each substrate, quadratic models were fit to the data. For the regolith + frass model (Eq. 3), the intercept (*p = 9.81E-13*) and quadratic (*p = 0.03124*) terms were both significant, and the linear term approached significance (*p = 0.0855*) suggesting a U-shaped curve. For the soil + frass model (Eq. 4), only the intercept was significant (*p = 2.65E-15*), with the quadratic (*p = 0.141*) and linear terms (*p = 0.122*) being insignificant, again suggesting co-linearity of terms or another non-linear relationship. The local maxima were found to be 20.03% and 14.24% for regolith and soil, respectively.

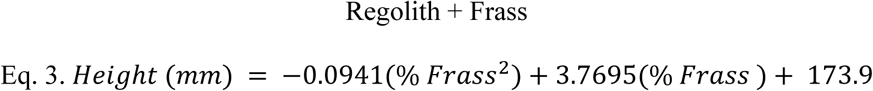

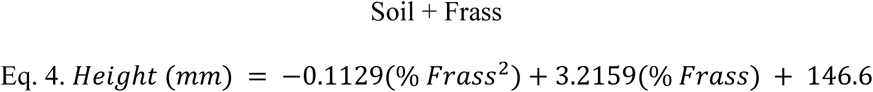

### 3.3 Chlorophyll Content

Leaf chlorophyll content was measured as an indicator of plant health [38]. Peas planted in regolith + frass generally exhibited higher chlorophyll levels at each tested concentration compared to those planted in soil + frass (Table 3). Notably, at 50% concentration no plants grew in soil, and so the mean chlorophyll content for peas planted in regolith + frass was obviously higher. Interestingly, a Pearson’s correlation test showed no significant correlation between leaf chlorophyll content and height for peas planted in regolith + frass (*R = -0.081, p = 0.68*), but that chlorophyll content was negatively correlated plant height (*R = - 0.48, p = 0.0095*) for peas planted in soil + frass. When pooling frass-inclusion levels, a Wilcoxon test showed a near-significant difference between the leaf chlorophyll levels between peas potted in regolith + frass and soil + frass (*p = 0.057*). When comparing leaf chlorophyll content in response to substrate type for the same levels of frass, Regolith and Frass were statistically different for 0%, 10%, and 50% frass inclusion, but not 25% (Table 3).

As above, regression models were fitted to the data to find optimal frass inclusion levels for each substrate. For plants potted in regolith + frass (Eq. 5), the intercept coefficient was highly significant (*p = 1.24E-13*), while the coefficients for both the linear (*p = 0.104*) and quadratic (*p = 0.230*) terms were not statistically significant, again suggesting co-linearity of terms or another non-linear relationship. For the soil + frass model (Eq. 6), neither the linear term (*p = 0.195*) nor the quadratic term (*p = 0.967*) were statistically significant, indicating that chlorophyll concentration did not vary meaningfully with frass addition within the tested range. The lower coefficients for the linear term and intercept in the regolith quadratic regression model compared to the soil model (Eq. 6) support the idea that chlorophyll levels were lower overall in peas potted in regolith.

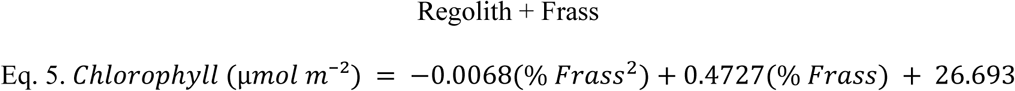

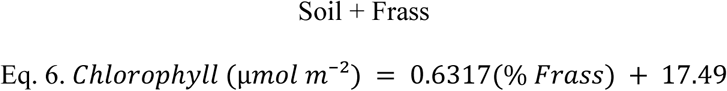

### 3.4 Biomass Production

Dry biomass weight of harvested fruit was used to evaluate overall plant productivity. A Mann-Whitney U test revealed a statistically significant difference in dry biomass between plants grown in regolith + frass compared to those grown in soil + frass (*U = 594.0, p = 0.0064*) (Figure 2). When comparing biomass production in response to substrate type for the same levels of frass, Regolith and Frass was statistically different for 0%, 10%, and 50% frass inclusion, but not 25% (Table 3). Importantly, no peas were produced when grown in soil + 50% frass.

**Figure 2.**
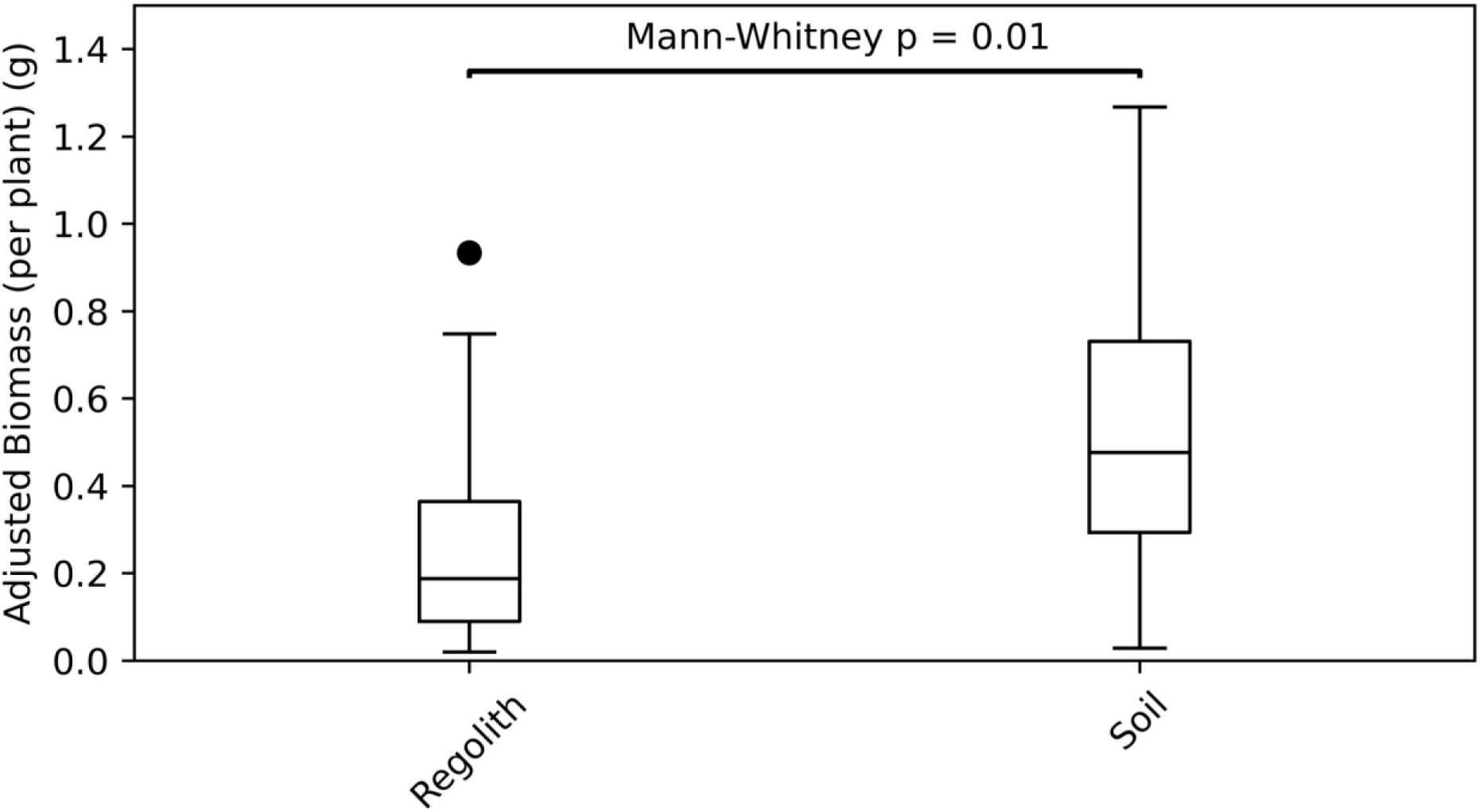
Boxplot comparing final biomass weight (g) between *Pisum sativum* grown in soil and regolith simulant. Each experimental unit consisted of two pea (Pisum sativum) seeds planted in 0.13-m tall, 0.15-m upper length & depth, and 0.105-m lower length & depth square pots filled with substrate, of which 0%, 10%, 25%, or 50% was BSF frass by volume. Mann-Whitney U test (ɑ = 0.05) was used to compare differences across treatments pooled by frass-inclusion percentage.

Once again, quadratic regressions were fit to the data to find optimal frass inclusion levels. For peas grown in regolith + frass (Eq. 7), the intercept was significant (*p = 0.001*), as was the linear (*p = 0.025*), and quadratic terms (*p = 0.020*), suggesting a u-shaped response. For peas grown in soil + frass, (Eq. 8) the intercept was significant (*p = 0.001*), as was the linear term (*p = 0.021*), but the quadratic term was not (*p = 0.071*), again suggesting co-linearity of another non-linear relationship. Still, quadratic regressions were chosen because visual analysis of the data indicated a single inflection point and non-constant trends, and these local maxima were found to be 22.13% and 19.62%, respectively.

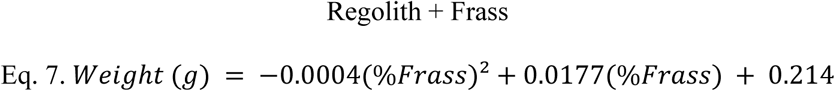

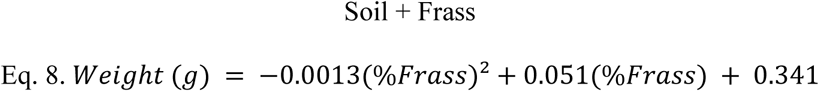

## 4. DISCUSSION

This study aimed to evaluate the growth and productivity of peas planted in Martian regolith simulant compared to those planted commercial gardening soil, both amended with increasing amounts of BSFL frass. During this study we assessed germination rates, plant height, chlorophyll content, and biomass production for peas grown in Martian regolith simulant and commercial gardening soil that were each combined with 0, 10, 25, and 50% BSFL frass by volume. The study found that germination rates were statistically similar between each substrate, and most interestingly, most peas (70.83%) germinated in regolith with 0% added frass, from which, 0.177 g (dry weight) of peas were harvested on average. Final plant height was also statistically similar between regolith and soil when pooling all frass inclusion levels together. Thus, the null hypothesis that peas could grow equally in Martian regolith and commercial gardening soil was generally supported, and this study also documents the possibility of growing peas in regolith without a frass amendment (though, the MMS-Simulant used here notably does not contain perchlorates).

When comparing the effect of frass inclusion on fitness for each treatment, notably, no plants survived the duration of the experiment when planted in soil + 50% frass, suggesting that high levels of frass can be toxic/detrimental to plant growth, especially if the soil is already nutrient rich. By contrast, small amounts of frass appear beneficial, since pea plants reached heights of 80.16 mm when planted in regolith + 10% frass, as compared to 41.46 mm in soil + 10% frass, a nearly 2-fold difference. Fitting quadratic models to the data revealed distinct optima of frass inclusion and for peas planted in regolith, our models suggest that pea germination rates will be optimized when supplementing with 6.9% frass, pea plant height will be maximized at 31.89% frass inclusion (although as discussed below, it may instead be better to minimize this quantity), and pea plant biomass production will be optimized at 22.19% frass inclusion; whereas these optima are 5.6%,14.24%, and 19.62% respectively. These different optima suggest not only a suite of life history trade-offs, but also a distinct plant physiological response to regolith.

### 4.1 Response in Regolith vs Commercial gardening soil

Interestingly, one of the primary differences we observed between peas planted in regolith vs soil treatments was the relationship between plant height and chlorophyll content. Peas planted in soil + frass showed a statistically significant negative correlation between height and chlorophyll content *(R = -0.48, p = 0.0095)*, indicating that taller plants tended to have lower chlorophyll levels. By contrast, peas planted in regolith simulant exhibited no such relationship *(R = -0.081, p = 0.68)*, suggesting that leaf tissues express consistent amounts of chlorophyll content regardless of how tall the pea plants grow. Since elevated chlorophyll may indicate improved nutrient uptake or reduced physiological stress [39], the opposite might be true as well, meaning comparatively less chlorophyll at high frass concentrations may indicate stress on peas planted in commercial potting mix. Our findings differ from past research on corn, *Zea mays,* and soybeans, *Glycine max,* which both showed a significant positive correlation between (*R = 0.87, p δ 0.001*) between chlorophyll content and plant height, though neither incorporated frass supplementation [38,40]. Considering this result indicates that perhaps peas grown in regolith + frass do not follow the same physiological pattern as do other plants potted in conventional soil.

Of course, since the optimal frass inclusion level for peas was 2.1-times higher when grown in regolith than in commercial gardening soil, this could potentially be due to differences in nutrition the peas received, or stress responses to being grown in Martian Regolith simulant, which plants obviously have not evolved to grow in. Although peas planted in regolith grew taller than their soil-planted counterparts, this may not necessarily be the best indicator for fruit production, since energy devoted to plant growth might trade off with fruit development. For example dwarf fruit trees mature much faster and some varieties produce more fruit than their conventional counterparts [41]. Future research is needed to understand such a possible trade-off occurring in peas when planted in Regolith.

Indications of plant stress were also seen in the ubiquitous increases in variation of fitness responses of plants grown in regolith compared to commercial gardening soil. For example, mean germination rates had a standard deviation of 39.65 in regolith + 0% frass, nearly double that the SD of 19.46 for peas grown in soil + 0% frass. Moreover, the SD for plant height in peas grown in regolith were also consistently ∼2-times greater than those grown in soil (Table 3), and the final biomass of peas grown in regolith + frass showed a higher coefficient of variation (i.e., ratio of the SD to the mean) across all frass inclusion levels. Considered holistically with our other findings, this higher variability suggests that future *in situ seedling* establishment in Martian regolith, while possible, will be erratic, since in many cases one or both of the seeds planted in each experimental unit were unsuccessful, but may also be circumvented by selectively breeding strains of peas that thrive best in regolith.

### 4.2 Alignment with Prior Work

The trends reported here are consistent with prior studies showing that excessive frass amendments can lead to nutrient imbalances, poor soil structure, or microbial shifts that negatively impact growth [32,42]. In our own pilot study, we found that peas cannot grow in substrates that contain more than 50% frass by volume, which likely represents the extreme limit of what can be tolerated by peas. Such high levels may contribute to nitrogen toxicity or other nutrient excesses, especially when added to commercial potting mix, and these excesses are known to induce physiological stress [43], such as through nitrogen ‘burn’ and an excess of salts which can damage and dehydrate roots, leading to stunted growth. Indeed, in some plants we observed yellowing and wilt (personal observation), though this could also have been due to high heat and/or sun-exposure within the greenhouse environment (even underneath shade cloth).

The results of this study also align with prior research on the use of BSFL frass as a biofertilizer. Previous studies on crops such as lettuce, *Lactuca sativa*, wild cabbage, *Brassica oleracea*, and tomatoes, *Solanum lycopersicum,* have demonstrated that frass amendments can enhance various characteristics of plant growth. For lettuce and tomatoes, 10 to 20% frass dosage by volume led to increased height, stem diameter, and leaf number over a 30-day period. For wild cabbage, an optimal frass dosage of ≃10% lead to increased plant weight and height [44]. Since the models presented in this study suggest an optimal range of frass inclusion between approximately 5-32%, taken together with these other studies, it shows that future research should explore a finer range of frass inclusion levels within this range.

### 4.3 Limitations

One major limitation of this study was that pea (fruit) was not harvested from pea pods, nor were the F1 generation planted again to test for viability. Instead, the total biomass of the plant was measured as a proxy for plant fitness when grown in Regolith. In addition, relatively large increments in frass-inclusion percentages tested (0%, 10%, 25%, 50%), may have obscured exactly where the ‘true’ optimal inclusion levels lie. To address this, we fit regressions to the data (and taking their derivatives yielded estimated optima), but these need to be verified experimentally, and future studies will need to confirm whether any peas grown in Regolith are (a) viable and (b) safe for human consumption.

Another limitation is that both trials of the experiment were conducted in a single greenhouse and were only separated in time by one week due to logistical constraints. Because of this overlap in both time and space, there may have been some level of pseudoreplication, since technically all the plants were grown within the same greenhouse environment [45]. However, differences in microenvironments from one side of the greenhouse to the other were likely significant enough to distinguish the trails from one another in terms of their ecology, since each experienced different lighting conditions (due to the east-west procession of the sun) temperature, and humidity (i.e., from being nearer or farther from the wet wall). While the greenhouse was equipped with cooling and ventilation systems, large temperature fluctuations did occur day-to-day due to the greenhouse windows being composed from untreated glass, which may have impacted the variation in plant growth, and because abiotic factors were measured at the spatiotemporal resolution of the entire greenhouse, they were not considered for the statistical analysis, but still included as part of supplementary data for future reference.

Lastly, a major consideration should be made regarding the Martian simulant used in this study, Mohave Mars Regolith Simulant (MMS-1), which is not a perfect replica of Marian Regolith because it is chiefly missing perchlorates, but has likewise been used by other studies [35]. As discussed, these are detrimental or toxic to plant growth, meaning the results of this study need to be considered lightly before making broad generalizations [46]. However, ongoing research is exploring the use of perchlorate-reducing bacteria and bioweathering to mitigate these toxic effects, which may enhance the viability of plant cultivation in Martian regolith in future applications [47]. Together these results point towards the possibility of future extraterrestrial farming.

## 5. CONCLUSION

In conclusion, this study showed that common garden peas can successfully be grown in Martian regolith simulant alone or when supplemented with BSFL frass, since statistically similar growth rates and biomass production were found to peas grown in commercial gardening soil. Fitting quadratic regressions to the data showed that the fitness of pea plants planted in regolith is predicted to be optimized at a frass inclusion level between 5-32%, depending on the metric being maximized. Together these results demonstrate that Martian regolith has the potential to support vegetable crop cultivation, marking a promising step towards *in situ* space agriculture. While past studies have considered horse/pig manure as an ISRU [48], this work furthers the viability of integrating nutrient cycling into bioregenerative life support systems from a much more feasible insect colony, since the study provides novel evidence of black soldier fly frass enhancing crop performance in a Martian regolith analog. These insights not only contribute meaningfully to astrobiology but can also act as a model to inform cyclical farming practices on Earth, such as in areas with degraded soil, considering growing crops in space takes place under an even more constrained system.

## ACKNOWLEDGEMENTS

The authors would like to thank their anonymous reviewers for help revising this manuscript, as well as The Martian Garden for providing Martian regolith simulants at a discounted price, as well as Andy Weir for the inspiration to pursue this work.

## AUTHOR VITAE

JEM - J. Emmanuel Mendoza is a Honors undergraduate student and Brown Scholar majoring in Aerospace Engineering at Texas A&M University. His research & development interests include spacecraft propulsion, agriculture for space applications, and human spaceflight. He has been a member of the F.L.I.E.S. Facility and the Aggie Sat Lab, helping pioneer a regolith simulant-frass plant growth project and create an autonomous LIDAR rover. He has interned for several Aerospace companies including Sierra Space and Axiom Space, and his work has appeared on *New York Science Times* and *National Public Radio*.

NBL - Noah Lemke earned his PhD in Entomology from Texas A & M University, during which he studied the reproductive behavioral ecology of the black soldier fly, *Hermetia illucens.* His research aims to link theory of insect science with application. His studies have been published in *BioEssays, Economic Entomology, Insects,* and *Journal of Insects as Food and Feed.* Presently, NBL is a Fulbright Scholar and Visiting Research Fellow at KU Leuven.

JKT - Dr. Jeffery K. Tomberlin is a Professor, AgriLife Research Fellow, & Presidential Impact Fellow in the Department of Entomology at Texas A&M University. Since arriving at Texas A&M University in 2002, 21 Ph.D. and 21 M.S. students have completed their degrees under his supervision. He has also supervised 8 postdoctoral associates. To date, he has edited 8 books, published 35 book chapters and +270 research articles which have more than 26,500+ citations. Through his efforts, he has been recognized as a Fellow by the American Academy of Forensic Sciences, Entomological Society of America, and the Royal Entomological Society.

## SUPPLEMENTARY INFO

**Supplemental Figure 1.**
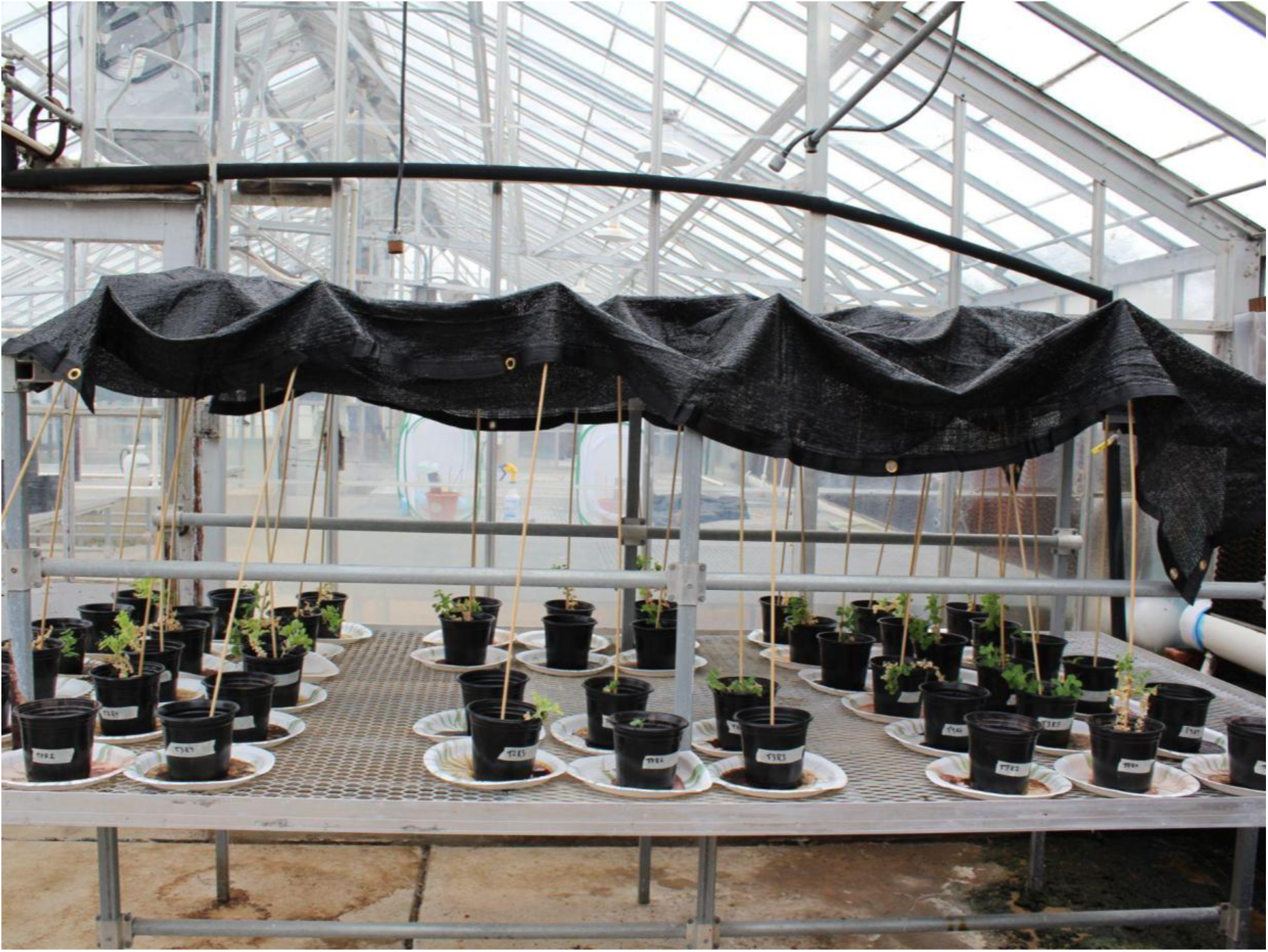
Experimental set-up for Trial A within the southwestern section of the FLIES lab greenhouse. Pea plants were randomly arranged in eight 2 × 3 arrays atop a rolling mesh-top bench. Each pot sat on a paper plate to prevent runoff, and 40% shade cloth was suspended 0.76 m above the bench to mitigate excess solar exposure.

## Notes

**Conflicts of Interest** Material used in this experiment was acquired from EVO Conversion Systems, LLC, a company which J K Tomberlin has significant financial interest in.

**Funding** This material is based upon work supported by the National Science Foundation under Grant Nos. 1746932, 2052565, 2052788, 2052454. Any opinion, findings, and conclusions or recommendations expressed in this material are those of the authors and do not necessarily reflect the views of the National Science Foundation.

### Competing Interest Statement

Material used in this experiment was acquired from EVO Conversion Systems, LLC, a company which J K Tomberlin has significant financial interest in.

